# KinLinks: Software Toolkit for Kinship Analysis and Pedigree Generation from HTS Datasets

**DOI:** 10.1101/046938

**Authors:** Anna Shcherbina, Darrell O. Ricke, Eric Schwoebel, Tara Boettcher, Christina Zook, Johanna Bobrow, Martha Petrovick, Edward Wack

## Abstract

The ability to predict familial relationships from source DNA in multiple samples has a number of forensic and medical applications. Kinship testing of suspect DNA profiles against relatives in a law enforcement database can provide valuable investigative leads, determination of familial relationships can inform immigration decisions, and remains identification can provide closure to families of missing individuals. The proliferation of High-Throughput Sequencing technologies allows for enhanced capabilities to accurately predict familial relationships to the third degree and beyond. KinLinks, developed by MIT Lincoln Laboratory, is a software tool that predicts pairwise relationships and reconstructs kinship pedigrees for multiple input samples using single-nucleotide polymorphism (SNP) profiles. The software has been trained and evaluated on a set of 175 subjects (30,450 pairwise relationships), consisting of three multi-generational families and 52 geographically diverse subjects. Though a panel of 5396 SNPs was selected for kinship prediction, KinLinks is highly modular, allowing for the substitution of expanded SNP panels and additional training models as sequencing capabilities continue to progress. KinLinks builds on the SNP-calling capabilities of Sherlocks Toolkit, and is fully integrated with the Sherlocks Toolkit pipeline.

## I. Introduction

Kinship testing may be used to identify relatives for immigration cases, or for intelligence in establishing relationships between individuals. For example, terrorist networks are often family centered [2]. Identifying relationships, both immediate and distant, may prove useful in better understanding networks at many levels. Kinship analysis may also enable genotype imputation to increase sample size in pedigreed populations for a number of secondary applications [3]. Kinship testing is needed for older remains identification with matching to second or third generation descendants. Yet another use involves deterring immigration fraud by verifying claimed familial relationships. For these purposes and others, the law enforcement community typically relies on sample identification by sizing of short tandem repeats (STRs) with capillary electrophoresis, usually coupled with searches against the CODIS database or a relative’s DNA. However, the ability of STRs to identify relatives is limited. Increasing the number of STR loci improves the statistical power of the analysis, but to a surprisingly modest degree [4]. Large panels of SNP loci may be useful in further resolving extended kinship relationships: chip-based assays using 192,000 loci can identify third-degree relationships [5]. One of the goals of this project was to achieve similar results with a smaller panel of loci detected by High-Throughput Sequencing (HTS).

A number of tools currently exist to identify first degree kinship relationships parents [6] [7]or siblings [8], but few tackle the more challenging problem of multi-generation pedigree reconstruction in the presence of missing or incomplete data. For example, the MPKin program allows users to select among multiple familial searching strategies to match a sample to a target database of potential relatives: minimum number of shared alleles, moderate stringency matches at all loci, Kinship Coefficient calculation, and other approaches [9]. However, though it has been used extensively by law enforcement agencies for pairwise relationship prediction, it stops short of full pedigree reconstruction.

Other tools go a step beyond pairwise kinship prediction to tackle the problem of automated pedigree reconstruction. IPED [10]and its successor IPEDX [11]use haplotype and identify by descent (IBD) information to reconstruct pedigrees generation-by-generation backwards in time. For each generation, the pairwise relationships are predicted between individuals within the current generation, and parents are created according to the predicted relationships. Though this approach is able to rapidly reconstruct pedigrees in the presence of perfect data and consanguineous marriages, it does not handle more complex relationships such as half-siblings or missing samples, both common phenomena for forensics analysis.

The KinLinks software combines some of the standard approaches to kinship prediction used by the above-mentioned tools, such as the Kinship Coefficient, probability of zero identity by state, and likelihood ratio calculation to identify the most likely pedigree from multiple candidates to predict pairwise kinship relationships and automatically reconstruct pedigrees for multiple SNP genotyped samples. KinLinks assumes no *a priori* knowledge about the relative ages of subjects, pedigree completeness (missing samples are expected and tolerated), or population allele frequencies. The algorithm training module uses a supervised machine learning classifier to build training models for pairwise kinship degree predictions. These pairwise predictions are then combined into multi-generation pedigrees via a set of heuristics to resolve relationship type (half siblings vs avuncular vs grandparent relationships for second degree relationships, for example) and direction (the parent in a parent/child relationship for example). Experimental validation on several hundred samples has yielded perfect prediction of first and second degree relationships as well as the ability to distinguish unrelated vs related individuals. Third degree relationships can be resolved with over 90% accuracy.

## II. Methods/Algorithm Overview

### A. SNP Panel Design

A panel of 5396 SNP loci was generated utilizing Ampliseq multiplex 150 bp amplicons. This amplicon size was compatible with the current capabilities of the Ion Torrent Proton instrument, which produces 100 million reads per run. These loci were chosen based on

- low minor allele frequency, based on the hypothesis that fractions of shared minor alleles are indicative of degree of relationship [14]
- low correlation with biogeographic ancestry, and
- maximal spacing along the genome (at least 50 kbp between loci)

### B. Training and test data samples

Source DNA from four groups was purchased from the Coriell Medical Institute to serve as training and test samples for kinship analysis. These include the Family 95 pedigree of 43 individuals across three generations, the ALS NINDS0760 family of 30 individuals across 4 generations, the Retinitis pigmentosa families 2110 and 2111 of 52 individuals across 6 generations (Figure1a), and 54 geographically diverse samples. SNP DNA from all individuals was analyzed using the custom-designed 5396-locus Ampliseq panel described above, with sequencing performed on the Ion Torrent Proton instrument. In phase 1 of the project, the pedigree generation algorithm was trained on 1225 pairwise relationships from Family 95 as well as 2862 pairwise relationships from the geographically diverse samples. The algorithm was tested on 870 pairwise relationships from the ALS family. In phase 2, the ALS samples were added to the training dataset, and algorithm performance was evaluated on the 2652 pairwise relationships within the Retinitis pigmentosa family.

**Fig. 1.**
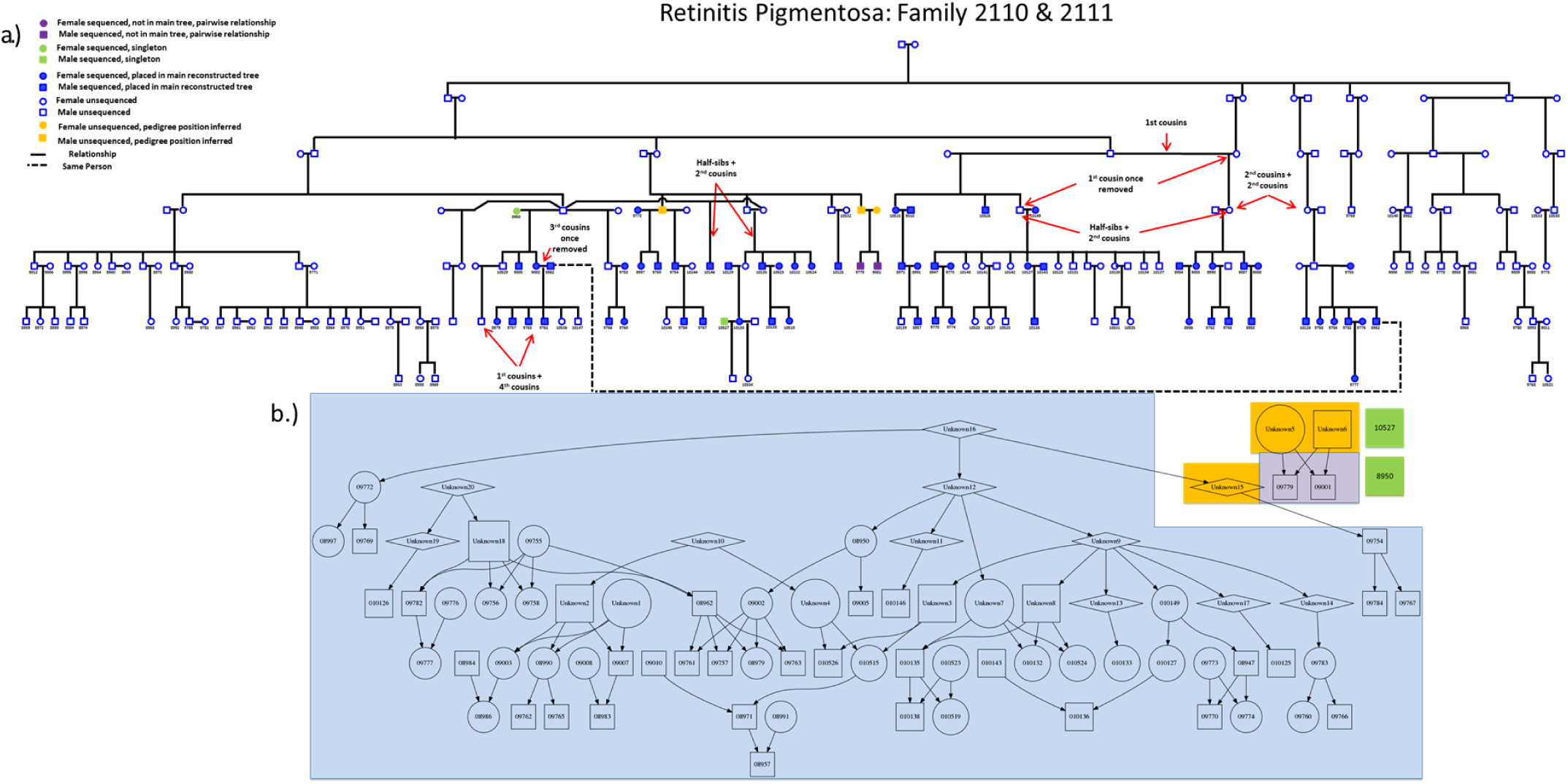
Automated pedigree generation for 52 members from the Retinitis pigmentosa family. a) Truth pedigree for the Retinitis pigmentosa family. b) Pedigrees for groups of sequenced profiles that were automatically generated by the kinship algorithm. Pedigrees are color-coded to correspond with truth data.

### C. Machine Learning Classifier for pairwise kinship prediction

Ten features were identified for training a one-vs-one multi-class support vector machine classifier. These include:

- probability of identity by state equaling zero P(IBS=0) [15], [16]. This metric measures the number of dissimilar alleles shared between a pair of individuals, where dissimilar is quantified as two individuals presenting with homozygous, opposite allele calls (one individual is homozygous minor, while the other is homozygous major). Unlike the Kinship Coefficient, this metric is able to differentiate parent-child relationships from siblings.
- Kinship Coefficient as calculated by the KING algorithm [17]. The Kinship Coefficient in this model is defined as the number of minor alleles shared between two individuals, divided by the average number of minor alleles between the two individuals. The Kinship Coefficient was selected because it is not reliant on population-estimated minor allele frequencies. In heterogeneous populations this type of estimation has been shown more accurate and robust to large groups of individuals [18], [19].
- biogeographic ancestry [18]predictions to the regional level (Americas, Europe, Middle East, East Asia, South Asia, East Africa, West Africa, Oceania) were determined via a genetic algorithm using a separate panel of 96 SNPs, as detailed in Ricke et al [1].
- number of shared loci.
- number of loci where both individuals were homologous for the minor allele.
- number of loci where both individuals were homozygous for the major allele.
- number of loci where both individuals where heterozygous for the minor allele.
- number of loci where one individual was homozygous for the minor allele while the other one was heterozygous.
- number of loci where one individual was homozygous for the major allele while the other was heterozygous.
- number of loci where one individual was homozygous for the major allele while the other was homozygous for the minor allele.

A one-vs-one support vector machine multi-class classifier with a linear kernel was implemented using the sklearn Python toolkit [20]. The classifier was trained via leave-one-out cross-validation on the set of pairwise relationships in Family 95 and the set of geographically diverse samples, as described above. Support vector machine parameters gamma and C were determined via a parameter sweep in two-dimensional space. Two classifiers were developed the first classifier predicts relationship degree (i.e. pairs of 1st, 2nd, 3rd degree relatives or unrelated individuals) and the second classifier predicts the exact relationship among a pair of samples (i.e. parent/child, sibling, grandparent, avuncular, cousin, unrelated). Both classifiers use the above-mentioned feature values for a set of samples as inputs. The first classifier predicts the degree of relation between each pair of samples and the second classifier predicts the relationship between each pair of samples.

### D. Heuristics for pedigree generation

An algorithm was implemented to automatically generate pedigrees from the pairwise kinship predictions. As a first step, a set of heuristics was implemented to determine relationship direction for first and second degree relatives (i.e. for a first-degree relationship, who are the parents, who are children, and who are siblings). These include:

- Spouse married into family: a sample has a first degree relationship to one or more other samples (children), but no relationship of any degree to other samples in the dataset.
- Siblings: siblings are expected to share 2 alleles at 25% of the loci, 1 allele at 50% of the loci, and 0 alleles at 25% of the loci.
- Direct descent: A child will share 50% alleles with a parent, 25% of alleles with a grandparent, 12.5% alleles with a great grandparent, 6.25% allele with a great grandparent.
- Trio pattern: If two unrelated samples both have a first degree relationship with a third sample, and each unrelated sample shares 50% of its alleles with the third sample, the first two samples are the parents of the third sample.

The python_graph tool library (v. 2.2.36) [27]was used to generate a semi-directed acyclic graph to represent the pairwise relationship predictions for a set of samples. Sex was inferred by analyzing sample genotypes at 30 loci present on the X chromosome. Additional heuristics were generated to infer connections between nuclear families, allowing the generation of fully-connected pedigrees for Coriell Family 95, the ALS family, and the Retinitis pigmentosa family (Figure1b). The auto-generated pedigrees were graphed as well as in family-tree format using the PyGraphViz toolkit (v.1.3rc2) [21].

Cousin relationships were resolved by identifying a unique pattern of allele sharing. All cousin relationships across the 72 individuals analyzed for Family 95 and ALS met the following criteria:

- Relationship identified as second or third degree by machine learning algorithm in the first step of kinship analysis.
- 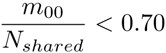
- 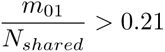
- 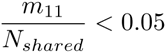

Nshared indicates the total number of loci in the kinship panel where both subjects had an allele call. *m*_00_ represents the total number of loci where both subjects had 0 minor alleles; *m*_01_ represents the number of loci where one subject had 0 minor alleles and the other had 1 minor allele; *m*_11_ represents the number of loci where both subjects had 1 minor allele and 1 major allele. If this pattern was observed between two nodes in the pedigree, the minimum number of nodes needed to create a cousin relationship were added to the pedigree. In cases where multiple assignments of nodes were possible, nodes were added in a way as to avoid any conflicts with existing high-confidence (nuclear family) relationships as well as to minimize conflicts with the degree designations of the machine learning algorithm in step 1. Some contradiction with the machine learning calls was tolerated, as the machine learning algorithm miscalls a fraction of relationships of degree 3 or higher.

Once cousins had been added to the graph, power calculations were performed to resolver relationships of the same degree. The purpose of this step was to determine whether each second degree relationship called by the machine learning algorithm was avuncular, grandparent-grandchild, or half-sibling. No difference in allele sharing patterns was observed between these three types of relationships, so the distinction was made by relying on graph connectivity and a modified form of the gradient ascent algorithm. The disjoint pedigree was treated as a set of connected components. The weighted degree of each node was computed, considering only nodes present in the other components (i.e. the nuclear families of which the node was not a member). Edges were weighted by the inverse of degree of relationship (i.e. a 2nd degree relationship was assigned w=1/2). This led to the observation that the weighted degree was higher for older individuals than for younger individuals in the same nuclear family, providing a means to infer age within nuclear families.

Furthermore, only individuals who had married into the family had a weighted degree of 0. The weightings also provide a set of constraints for resolution of second degree relationships. For example, grandparents will have degree weightings three to four times higher than their grandchildren. Half siblings will have similar degree weightings if the shared parent is part of the pedigree. If the shared parent is not part of the pedigree, one of the siblings will have a degree weight of 0, while the other will have a non-zero degree weight. This gives rise to the following set of constraints:

- If two nodes have similar degree weightings, they do not share a grandparent/grandchild relationship.
- If one node has a degree weight of 0, while the other has a non-zero degree weight, they are half-siblings related through a parent that married into the family.
- If two nodes have degree weightings of ratio greater than 2, but neither weighting is 0, they are not half-siblings.
- If two nodes share a grandparent/grandchild relationship, the grandparent node will have a higher degree weight.

Given this set of constraints, a set of second degree relationships were selected to avoid contradictions with established nuclear family relationships and minimize contradictions with the machine learning algorithm degree calls from step one. For example, if node A and B have similar degree weightings, they may be either half-siblings or have an avuncular relationship. The minimum possible number of nodes is added to the pedigree to support each relationship and any contradictions are computed. The relationship that minimizes contradictions is assigned. In case of ties, a relationship is not assigned for the two nodes. All second degree relationships are examined, and the algorithm is repeated until no additional second degree relationships can be resolved.

### E. Resource Requirements

KinLinks is implemented in Python (version 2.7), and was evaluated on a machine running Fedora 7. The training phase of the algorithm was executed in 43 minutes, when training on a set of 124 samples (15252 pairwise relationships) using a single core and 8 GB of RAM. A training model can be generated once and stored for repeated use with test samples. This allows the test phase of the algorithm to be executed in under 1 minute for an equal number of relationships.

## III. Results

Supplementing the traditional Kinship Coefficient and P(IBS=0) metrics with the other 8 features used for the one-vs-one machine learning classifier yields improved results, as presented in Tables I, II. Table II illustrates the performance of KinLinks when the algorithm is trained on both the Family 95 and ALS datasets, along with 54 geographically diverse unrelated samples, and tested on the Retinitis pigmentosa family. All 40 parent-child relationships and 22 sibling relationships were correctly identified correctly. For second degree relationship predictions, 77% accuracy was achieved. The remaining 23% of individuals were classified to within 1 degree of relation (either 1st or 3rd degree relatives). Though 56 individuals were correctly classified as second degree relatives, KinLinks had more difficulty in distinguishing between specific second degree relationships - grandparents, versus aunts/uncles, versus half siblings. Accuracy falls to below 50% in classifying relationships of degree four and higher. 94% of unrelated individuals are correctly identified as such. Overall, on the Retinitis pigmentosa test family, KinLinks has a false positive rate of 6% and an overall false negative rate of 24%, where the false negative rate is the defined as the failure to identify a relationship among two individuals with relationship of degree 5 or lower.

**Table I.** Degree relationship confusion matrix for 52 samples from the retinitis pigmentosa family. The predictions were made with a classifier trained on Family 95, ALS, and 54 unrelated samples of diverse biogeographic backgrounds. Relationships that were predicted correctly are highlighted in green, predictions that differed by 1 degree from the truth are highlighted in yellow, predictions that differed from the truth by 2 or more degrees are highlighted in red.

|  |  | Truth |  |  |  |  |  |  |  |  |  |  | Unrelated |
| --- | --- | --- | --- | --- | --- | --- | --- | --- | --- | --- | --- | --- | --- |
|  |  | 1 | 2 | 3 | 4 | 5 | 6 | 7 | 8 | 9 | 10 | 11 |  |
| Predicted | 1 | 63 | 0 | 0 | 0 | 0 | 0 | 0 | 0 | 0 | 0 | 0 | 0 |
|  | 2 | 0 | 56 | 4 | 0 | 1 | 0 | 0 | 0 | 0 | 0 | 0 | 4 |
|  | 3 | 0 | 18 | 41 | 20 | 14 | 9 | 0 | 3 | 0 | 2 | 0 | 14 |
|  | 4 | 0 | 0 | 4 | 13 | 9 | 12 | 6 | 3 | 0 | 4 | 0 | 23 |
|  | 5 | 0 | 0 | 0 | 2 | 1 | 3 | 2 | 2 | 0 | 3 | 1 | 6 |
|  | 6 | 0 | 0 | 0 | 0 | 0 | 0 | 0 | 0 | 0 | 1 | 0 | 1 |
|  | 7 | 0 | 0 | 0 | 0 | 0 | 0 | 0 | 0 | 0 | 0 | 0 | 0 |
|  | 8 | 0 | 0 | 0 | 0 | 0 | 0 | 0 | 0 | 0 | 0 | 0 | 0 |
|  | 9 | 0 | 0 | 0 | 0 | 0 | 0 | 0 | 0 | 0 | 0 | 0 | 0 |
|  | 10 | 0 | 0 | 0 | 0 | 0 | 0 | 0 | 0 | 0 | 0 | 0 | 0 |
|  | 11 | 0 | 0 | 0 | 0 | 0 | 0 | 0 | 0 | 0 | 0 | 0 | 0 |
|  | Unrelated | 0 | 0 | 1 | 16 | 59 | 111 | 58 | 62 | 1 | 58 | 10 | 657 |

**Table II.**
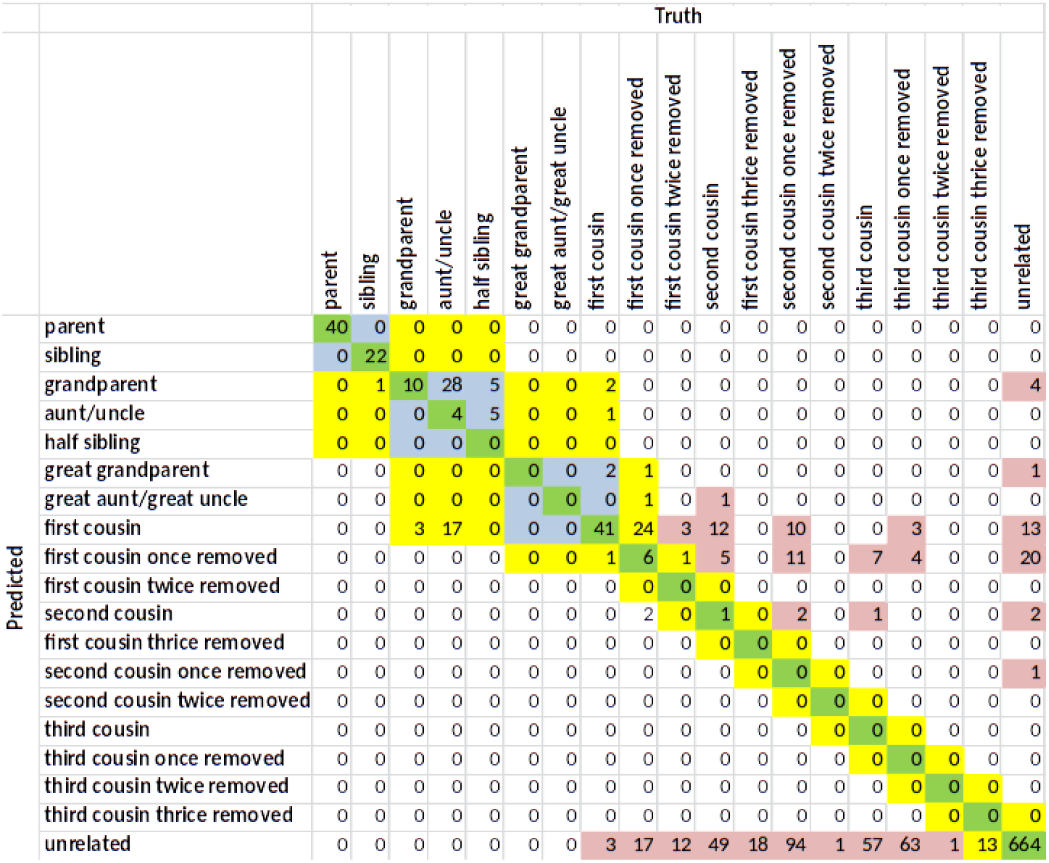
Confusion matrix for 52 samples from the Retinitis pigmentosa family; exact relationship PREDICTIONS. Predicted relationships that differ from the truth but share the same degree of relation are highlighted in blue.

The Retinitis pigmentosa family presents two challenges for classification. The first is the presence of consanguineous marriages [22], annotated by red arrows in Figure 1a. Consequently, a number of relationships do not fit the training model generated by Family 95 relationships as well as the ALS family. For the ALS family, no cases were observed where the predicted degree of a relationship was more than 1 degree closer than the truth degree. However, for the Retinitis pigmentosa family, 1.8 of predictions exhibit this error. In the most extreme cases, 4 pairs of unrelated individuals were predicted as second degree relatives, and 9 pairs were predicted as third degree relatives. A second challenge can be attributed to the high proportion of non-sequenced samples (non-highlighted nodes in Figure 1a) in the Retinitis pigmentosa family. Consequently, four sub-pedigrees are generated for the Retinitis pigmentosa family (Figure 1b). KinLinks is able to interpolate missing nodes, denoted by Unknown labels, that connect second degree relatives and cousins, up to 4th degree relationships.

The influence of biogeographic ancestry on kinship prediction was examined (Figure 2). In the first KinLinks iteration, biogeographic ancestry predictions were not included as a feature for the one-vs-one SVM classifier. When the algorithm was trained on Family 95, of European descent, as well as 54 geographically diverse individuals, a number of errors were present in the relationship predictions (Figure 2a). Unrelated individuals from the same world region frequently showed up as having a second, third, or fourth degree relationship. A cluster of 6 South American subjects (purple nodes) were predicted as related to each other, as was a cluster of five European subjects. Subsequently, a genetic algorithm was implemented to predicted sample biogeographic ancestry to major world region [1]. A test of the algorithm on the ALS family in conjunction with the 54 geographically diverse samples removed all relationships between unrelated pairs of individuals, with the exception of two (Figure 2b). One remaining second degree relationship was between a pair of Quechua individuals, and the second remaining third degree relationship was between a pair of Druze samples. Since relationship annotations used to generate the truth data are self-reported, it could not be established whether these are indeed false positives or unreported true positive relationships.

**Fig. 2.**
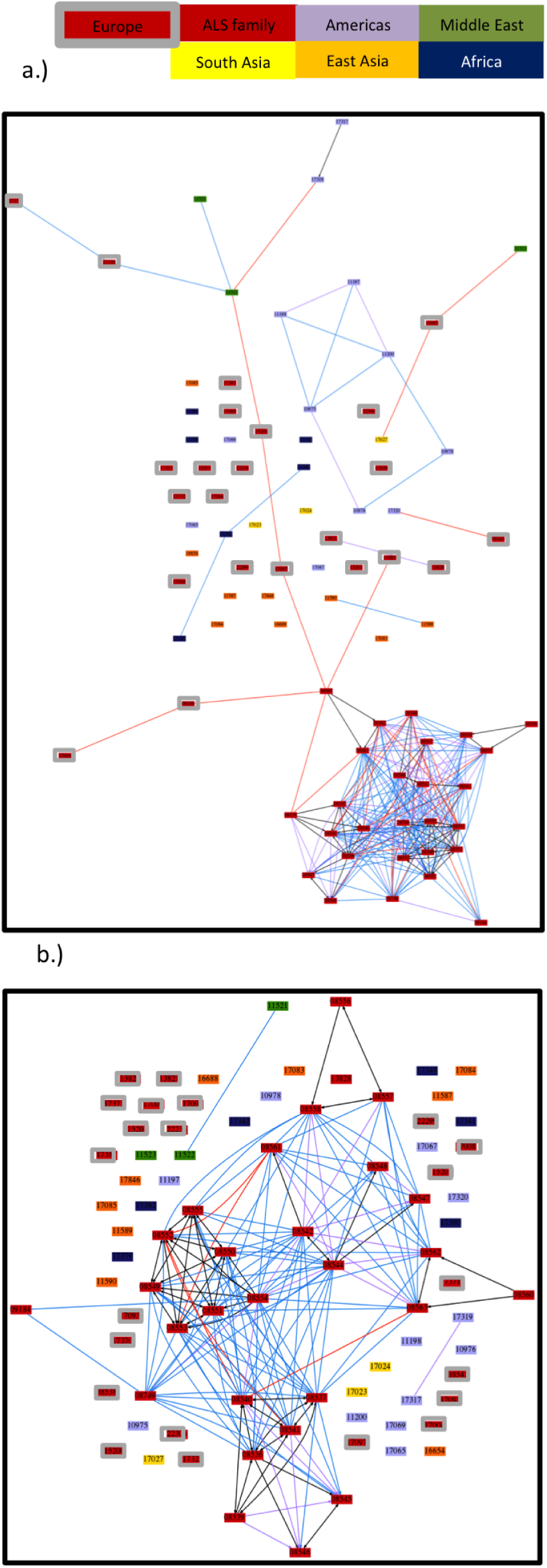
Effect of biogeographic ancestry on classifier performance. Predictions were made for 29 samples from the ALS family (red nodes), 17 unrelated European samples (red nodes with gray outline), 12 Central/South American samples (purple nodes), three Middle Eastern samples (green nodes), 9 East Asian samples (orange nodes), three South Asian samples (yellow nodes), 6 African samples (dark blue nodes). a) Relationship predictions when biogeographic ancestry was not utilized. b) Relationship predictions after biogeographic ancestry was added to the classification algorithm.

## IV. Discussion

The current panel size of 5396 SNPs is sufficient to resolve first degree relationships with 100% and second degree with 75% percent accuracy. Unrelated individuals can also be identified with 94% accuracy, and third degree relatives can be called with over 80% accuracy. However, the algorithm is unable to resolve relationships of degree 4 or higher with accuracy over 50% [23]. Algorithm performance was evaluated for varying SNP panel sizes, ranging from a 674 SNP subset of the panel (1/8 of the SNPs) to the full 5396 SNPs. By fitting three-dimensional splines to the performance curves, performance for larger SNP panels can be extrapolated. This method suggests that doubling the panel size to 10792 SNPs will enable resolution of 4th degree relationships to within 1 degree of relatedness. Tripling the panel size to 15000 SNPs is likely to enable perfect resolution of 4th degree relationships. However, the extrapolation suggests that a panel size of 690,000 SNPs would be necessary to resolve 5th degree relationships to within 1 degree. Improving SNP panel design provides an alternative to increasing panel size for resolution of higher degree relationships. Analysis of linked SNPs [24]or shared chromosome segments [25]has the potential to increase power of resolution. Advances in microhaplotype analysis techniques enabled by NGS sequencing may be useful for resolving higher degree relationships as well [26]. Furthermore, the current panel includes 30 SNPs from the X chromosome, and no SNPs from the Y chromosome. By including a sub-panel of X and Y SNPs in the analysis, patterns of X/Y inheritance can be traced through a pedigree and used to generate additional features for raining the one-vs-one SVM classifier. Work has been initiated on a second iteration of KinLinks, with a second panel of X and Y SNPs that will be used to resolve relationships within a predicted degree (i.e. grandparent vs avuncular). Thirdly, as the cost of exome and whole genome sequencing continues to drop, KinLinks can be modified to work with whole genome sequence data [27].

## REFERENCES

[1] Ricke D, Shcherbina A, Chiu N, Harper J, Petrovick M, Boettcher T, et al. Sherlock’s Toolkit: A Forensic DNA Analysis System. arXiv. 2015.

[2] Magouirk J, Atran S, Sageman M. Connecting Terrorist Networks. Studies in Conflict & Terrorism. 2008;31.

[3] Hickey JM, Cleveland MA, Maltecca C, Gorjanc G, Gredler B, Kranis A. Genotype imputation to increase sample size in pedigreed populations. Methods Mol Biol. 2013;1019:395–410.

[4] O’Connor K, Butts E, Hill C, Butler J, Vallone P. Evaluating the Effect of Additional Forensic Loci on Likelihood Ratio Values for Complex Kinship Analysis. The Twenty-First International Symposium Madioson, WI: Promega; 2010.

[5] Keating B, Bansal A, Walsh S, Millman J, Newman J, Kidd K, et al. First all-in-one diagnostic tool for DNA intelligence: genome-wide inference of biogeographic ancestry, appearance, relatedness, and sex with the Identitas v1 Forensic Chip. Int J Legal Med. 2013;127:559–72.

[6] Hayes BJ. Efficient parentage assignment and pedigree reconstruction with dense single nucleotide polymorphism data. J Dairy Sci. 2011;94:2114–7.

[7] Anderson EC. Large-scale parentage inference with SNPs: an efficient algorithm for statistical confidence of parent pair allocations. Stat Appl Genet Mol Biol. 2012;11.

[8] Ashley MV, Caballero IC, Chaovalitwongse W, Dasgupta B, Govindan P, Sheikh SI, et al. KINALYZER, a computer program for reconstructing sibling groups. Moleculary Ecology Resources. 2009;9:1127–31.

[9] Ge J, Budowle B, Chakraborty R. DNA identification by pedigree likelihood ratio accommodating population substructure and mutations. Investigative Genetics. 2010;1.

[10] He D, Wang Z, Han B, Parida L, Eskin E. IPED: Inheritance Path-based Pedigree Reconstruction Algorithm Using Genotype Data. Journal of Computational Biology. 2013;20:780–91.

[11] He D, Eskin E. IPEDX: An exact algorithm for pedigree reconstruction using genotype data. Bioinformatics and Biomedicine (BIBM), 2013 IEEE International Conference on2013. p. 517–20.

[12] Kling D, Tillmar AO, Egeland T. Familias 3 Extensions and new functionality. Forensic Science International: Genetics. 2014;13:121–7.

[13] Riester M, Stadler PF, Klemm K. FRANz: reconstruction of wild multi-generation pedigrees. Bioinformatics. 2009;25:2134–9.

[14] Ma W, Yang Y, Chen ZZ, Wang L. Mutation region detection for closely related individuals without a known pedigree using high-density genotype data. IEEE/ACM Trans Comput Biol Bioinform. 2012;9:499–510.

[15] Kirkpatrick B, Li SC, Karp RM, Halperin E. Pedigree reconstruction using identity by descent. Journal of Computational Biology. 2011;18:1481–93.

[16] Huff CD, Witherspoon DJ, Simonson TS, Xing J, Watkins WS, Zhang Y, et al. Maximum-likelihood estimation of recent shared ancestry (ERSA). Genome Res. 2011;21:768–74.

[17] Manichaikul A, Mychaleckyj J, Rich S, Daly K, Sale M, Chen W. Robust relationship inference in genome-wide association studies. Bioinformatics. 2010;26.

[18] Amirisetty S, Hershey GKK, Baye TM. AncestrySNPminer: A bioinformatics tool to retrieve and develop ancestry informative SNP panels. Genomics. 2012;100:57–63.

[19] Boukaze C, Keyser C, Crubezy E, Montagnon D, Ludes B. Pigment phenotype and biogeographic ancesty from ancient skeleatl remains: inferences from multiplexed autosomal SNP analysis. Int J Legal Med. 2009;123:315–25.

[20] Pedregosa F, Varoquaux G, Gramfort A, Michel V, Thirion B, Grisel O, et al. Scikit-learn: Machine Learning in Python. Journal of Machine Learning Research. 2011;12:2825–30.

[21] Hagberg A. PyGraphviz. 2014.

[22] Liu EY, Zhang Q, McMillan L, de Villena FPM, Wang W. Efficient genome ancestry inference in complex pedigrees with inbreeding. Bioinformatics. 2010;26:i199–i207.

[23] Egeland T, Pinto N, Vigeland MD. A general approach to power calculation for relationship testing. Forensic Science International: Genetics. 2014;9:186–90.

[24] Silberstein M, Weissbrod O, Otten L, Tzemach A, Anisenia A, Shtark O, et al. A system for exact and approximate genetic linkage analysis of SNP data in large pedigrees. Bioinformatics. 2013;29:197–205.

[25] Axenovich TI, Aulchenko YS. MQScore SNP software for multipoint parametric linkage analysis of quantitative traits in large pedigrees. Ann Human Genet. 2010;74:286–9.

[26] Kidd KK, Pakstis AJ, Speed WC, Lagace R, Chang J, Wootton S, et al. Current sequencing technology makes microhaplotypes a powerful new type of genetic marker for forensics. Forensic Science International: Genetics. 2014;12.

[27] Li H, Glusman G, Hu H, Shankaracharya J, Caballero J, Hubley R, et al. Relationship estimation from whole-genome sequence data. PLoS Genet. 2014;10:e1004144.

